# Candidate cancer driver mutations in superenhancers and long-range chromatin interaction networks

**DOI:** 10.1101/236802

**Authors:** Lina Wadi, Liis Uusküla-Reimand, Keren Isaev, Shimin Shuai, Vincent Huang, Minggao Liang, J. Drew Thompson, Yao Li, Luyao Ruan, Marta Paczkowska, Michal Krassowski, Irakli Dzneladze, Ken Kron, Alexander Murison, Parisa Mazrooei, Robert G. Bristow, Jared T. Simpson, Mathieu Lupien, Michael D. Wilson, Lincoln D. Stein, Paul C. Boutros, Jüri Reimand

## Abstract

A comprehensive catalogue of the mutations that drive tumorigenesis and progression is essential to understanding tumor biology and developing therapies. Protein-coding driver mutations have been well-characterized by large exome-sequencing studies, however many tumors have no mutations in protein-coding driver genes. Non-coding mutations are thought to explain many of these cases, however few non-coding drivers besides *TERT* promoter are known. To fill this gap, we analyzed 150,000 *cis*-regulatory regions in 1,844 whole cancer genomes from the ICGC-TCGA PCAWG project. Using our new method, ActiveDriverWGS, we found 41 frequently mutated regulatory elements (FMREs) enriched in non-coding SNVs and indels (*FDR*<0.05) characterized by aging-associated mutation signatures and frequent structural variants. Most FMREs are distal from genes, reported here for the first time and also recovered by additional driver discovery methods. FMREs were enriched in super-enhancers, H3K27ac enhancer marks of primary tumors and long-range chromatin interactions, suggesting that the mutations drive cancer by distally controlling gene expression through threedimensional genome organization. In support of this hypothesis, the chromatin interaction network of FMREs and target genes revealed associations of mutations and differential gene expression of known and novel cancer genes (*e.g., CNNB1IP1, RCC1*), activation of immune response pathways and altered enhancer marks. Thus distal genomic regions may include additional, infrequently mutated drivers that act on target genes via chromatin loops. Our study is an important step towards finding such regulatory regions and deciphering the somatic mutation landscape of the non-coding genome.

## Introduction

Cancer is driven by somatic driver mutations such as single nucleotide variants (SNVs), insertions-deletions (indels) and copy number alterations (CNAs) that affect critical genes and pathways. Driver mutations unlock oncogenic cellular properties of unconstrained proliferation, replicative immortality, immune evasion and the other hallmarks of cancer^1^. Completing the catalogue of cancer driver mutations is a central challenge of cancer research and key to understanding tumor biology, developing precision therapies and molecular biomarkers.

The search for driver mutations is complicated by the high rate of somatic ‘passenger’ mutations that have no biological significance. Statistical methods are used to distinguish between drivers and passengers in cancer genome sequencing datasets. These methods assume that somatic driver mutations occur more frequently than expected from background mutation rates, have unexpectedly high functional impact and show enrichment in biological pathways and networks (reviewed in ^2–4^). Driver discovery is facilitated by large genomic datasets assembled by consortia like the International Cancer Genome Consortium (ICGC)^5^ and The Cancer Genome Atlas (TCGA)^6^. The notable driver mutation in the *TERT* promoter that confers replicative immortality on cells by inhibiting telomere-related cellular senescence was first identified in melanoma^7,8^ and then in pan-cancer analyses^9,10^. These mutations create new transcription factor (TF) binding sites (TFBS) which increase *TERT* transcription^11^. Other genes with frequent promoter mutations include the protein-coding genes *PLEKHS1, WDR74* and SDHD^9–10^ along with the long non-coding RNAs (lncRNAs) *NEAT1* and *MALAT1*^12^. Genome-wide driver discovery studies are limited to gene-focused genomic regions such as promoters and untranslated regions (UTRs) rather than experimentally defined regulatory regions. Alternative approaches have scanned the genome with fixed-width windows^10,13^, defined windows around mutation hotspots^9,14^, or annotated cancer mutations using *cis*-regulatory information^14–15^. Window-based approaches do not capture the precise boundaries of regulatory elements while annotation-based approaches conduct limited statistical testing of mutations. Current approaches are also unable to determine potential target genes of distal mutations.

Driver discovery in the non-coding regulatory genome is challenged by complex overall distribution of somatic mutations. At the megabase scale, mutation burden is associated with transcriptional activity and replication timing^16,17^. Open chromatin is generally characterized by fewer somatic mutations while enhancers of the tissue of origin accumulate more mutations^18,19^. At the nucleotide scale, mutation signatures are manifested in uneven distribution of mutations in their trinucleotide context. Different signatures are characteristic of different tumor types and have been linked to aberrant activity of DNA repair pathways, effects of various carcinogens or molecular clocks^20^. Genome-wide analyses of short sequence motifs bound by TFs have revealed increased mutation rates in regulatory regions^21^, for example excessive promoter mutations melanoma and other cancer types are likely explained by decreased activity of the nucleotide excision repair pathway^22,23^. These studies suggest that a large fraction of gene regulatory mutations are caused by local mutational processes rather than positive selection driving tumor evolution.

The eukaryotic genome is organized three-dimensionally in the nuclear space to enable its functions, including transcription regulation via long-range interactions of promoters and enhancers and TF binding^24^. Binding sites of the CTCF chromatin architectural factor and the cohesin complex subunit RAD21 co-occur at topologically associated domain boundaries engaged in long-range chromatin interactions^24,25^ and are frequently mutated in colorectal cancer^26^. Anchors of chromatin interactions include functional genetic polymorphisms^27,28^ and are enriched in mutations in liver and esophageal cancers^29^. The *MYC* super-enhancer locus at 8q24 harbors SNVs with genetic predisposition for multiple tumor types^30,31^ and its deletion in mice was recently associated with reduced tumorigenesis^32^. Recurrent somatic mutations in enhancers of *PAX5* and *TAL1* have been found in leukemia and associated with differential gene expression^33,34^. Structural rearrangements in medulloblastoma and leukemia cause enhancer hijacking where oncogene expression is induced through translocations that associate oncogenes with active enhancers^35,36^. Thus some mutations at gene regulatory sites may drive cancer by re-configuring gene regulatory interactions or the three-dimensional folding of chromatin. Surprisingly, then, a systematic driver analysis of non-coding mutations in c/s-regulatory and three-dimensional chromatin interaction networks is currently lacking.

To fill this gap and to explore the effects of non-coding somatic mutations on gene-regulatory networks, we used 2,583 tumor-normal pairs characterized with whole genome sequencing (WGS) by the ICGC-TCGA Pan-cancer Analysis of Whole Genomes (PCAWG) project. We identified candidate drivers in regulatory regions of the human genome defined by the Encyclopedia of DNA Elements (ENCODE)^37^, then integrated these with the three-dimensional architecture of the human genome to prioritize and interpret candidate non-coding cancer drivers and their potential target genes. We found dozens of frequently mutated regulatory elements (FMREs) that were enriched in somatic small mutations and structural variants and overrepresented in active regulatory elements. Mutations in FMREs associated with altered expression of target genes, suggesting that our findings include novel driver mutations that rewire gene regulatory networks.

## Results

### Genome-wide discovery of cancer driver mutations with ActiveDriverWGS

We used the ICGC-PCAWG dataset of 2,583 whole cancer genomes for driver discovery and focused on mutations from 1,844 genomes from 31 cancer types, comprising 14.2 million single nucleotide variants and *indels^[PCAWG marker paper]^* (**Supplementary Figure 1**). We excluded four cancer types with atypical mutational processes: melanomas with elevated mutation rates in active TFBS^22^, lymphomas with localized hypermutations^38^, and liver and esophageal cancers with frequent mutations in topologically associated CTCF binding sites^29^ (**Supplementary Note, Supplementary Figure 2**). We also excluded a small subset of hypermutated tumors (69) that carried 47% of all somatic mutations.

To find non-coding cancer drivers in whole cancer genomes, we created ActiveDriverWGS, a genome-wide driver discovery method that statistically identifies genomic regions with an elevated frequency of somatic mutations (**Figure 1a**). ActiveDriverWGS performs a statistical analysis of single nucleotide variants (SNVs) and small insertions-deletions (indels) relative to adjacent background sequences using Poisson generalized linear regression, expanding our earlier work on protein-coding drivers^39^. The model estimates expected mutation burden through a relatively narrow adjacent window and is therefore less sensitive to mega-base scale fluctuation of mutation rates. To adjust for nucleotide-level mutational signatures that vary considerably across patients and tumor types^20^, the model includes covariates for the frequency of each mutation type in its trinucleotide context. ActiveDriverWGS additionally predicts mutation impact by detecting frequently mutated binding sites within candidate driver genes and noncoding regions.

**Figure 1.**
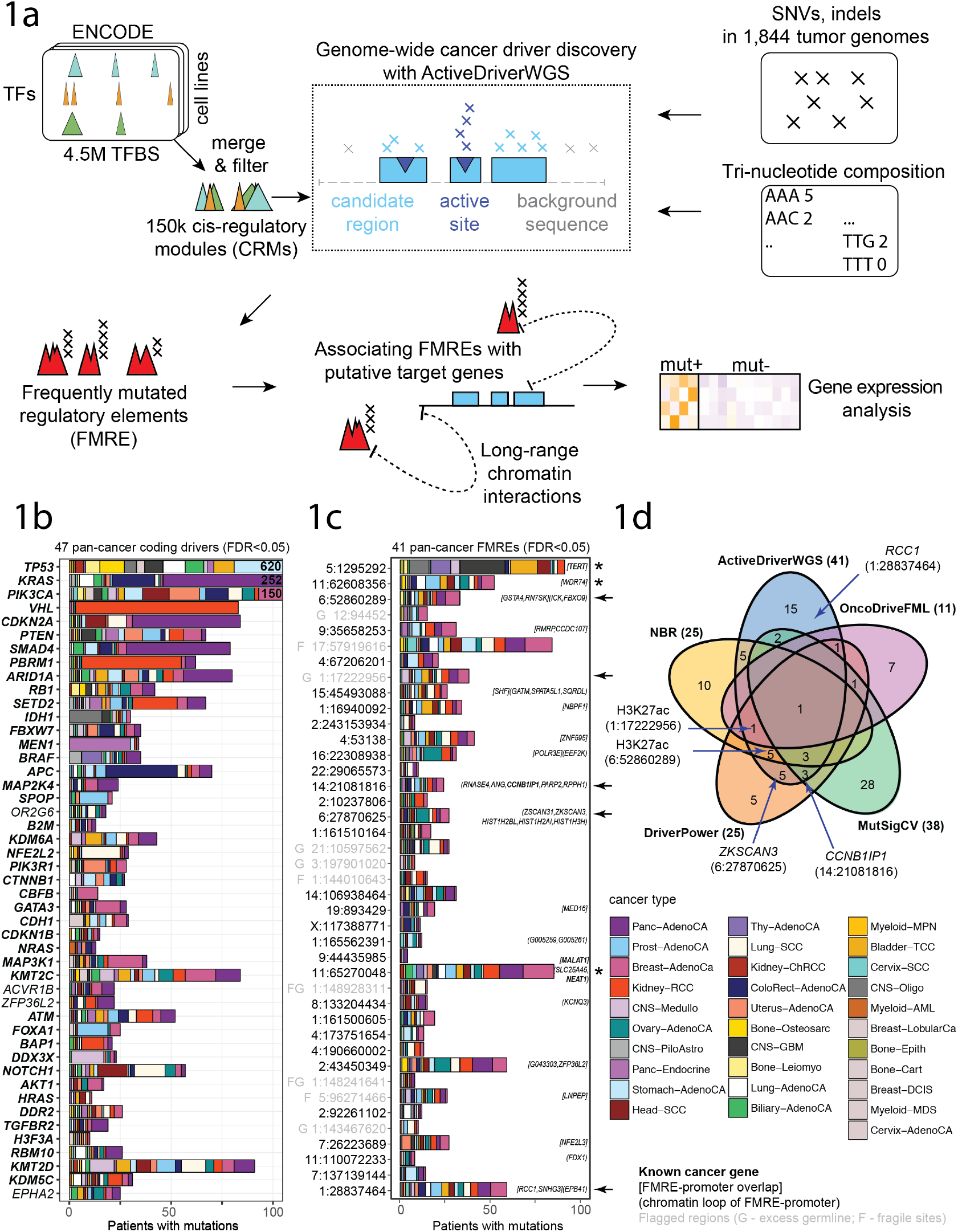
Cancer driver discovery in regulatory regions of the genome. (**a**) Discovery of frequently mutated regulatory elements (FMREs) as candidate cancer drivers. We analyzed cis-regulatory modules (CRMs) comprising clustered transcription factor binding sites (TFBS) from ChIP-seq datasets in ENCODE that were conserved in multiple cell lines and bound by at least two TFs. Single nucleotide variants (SNVs) and small indels from the PCAWG WGS dataset were used for driver discovery. Our novel genome-wide driver discovery method ActiveDriverWGS evaluates the enrichment mutations in candidate driver regions relative to adjacent background sequence and trinucleotide sequence content. Candidate non-coding drivers (FMREs) were then associated to potential target genes using long-range chromatin interactions derived from public HiC datasets. To validate candidate drivers, we associated FMRE mutations with gene expression changes of target genes. (**b**) Protein-coding drivers detected in analysis of the pancancer cohort. Known cancer drivers annotated in the Cancer Gene Census database are printed in bold. (**c**) Frequently mutated regulatory elements (FMREs) detected in pan-cancer analysis of CRMs. Genes associated with FMREs are shown right of the bars. Arrows show FMREs highlighted in the manuscript and asterisks indicate previously known non-coding driver regions. FMREs with gray labels are flagged due to excess germline variation in PCAWG. (**d**) Comparison of FMREs identified by five driver discovery methods. Two thirds of FMREs identified by ActiveDriverWGS are also found by at least one other method.

We validated ActiveDriverWGS by confirming its ability to recover known protein-coding and non-coding cancer drivers in the pan-cancer cohort and individual cancer types. We detected 47 coding genes (*FDR*<0.05) in a pan-cancer analysis, including 43 known drivers annotated in the Cancer Gene Census database^40^ (Fisher’s exact *P*=3.0×10^−62^, **Figure 1b**). Driver analyses of 31 cancer type specific cohorts revealed 70 genes and 59 known drivers in total (**Supplementary Figure 3**). Among non-coding consensus regions studied in *PCAWG^[PCAWG-2-5-9-14]^*, we recovered previously described non-coding regions with frequent mutations such as promoters of *TERT* and *WDR74*, the lncRNAs *NEAT1* and *MALAT1* as well as other candidates (**Supplementary Figure 4**).

We benchmarked ActiveDriverWGS and found that our statistical framework is well-calibrated. We tested three independently generated sets of simulated somatic mutations including two from the PCAWG project and one internally generated set (**Supplementary Figure 5**). We also tested three configuration changes in the driver discovery pipeline: genomic window sizes for determining background mutations, inclusion of hyper-mutated samples, and exclusion of model cofactors corresponding to trinucleotide sequence composition. ActiveDriverWGS was robust to the size of the background window, and our simulations showed that statistical strength was maximized with a 50 kbp window size. We further confirmed the importance of using trinucleotides for driver discovery, as exclusion of this cofactor greatly increased false positive findings among protein-coding drivers (47 vs 4 non-cancer genes found). As anticipated, inclusion of hypermutated samples in the pan-cancer analysis led to recovery of fewer known protein-coding drivers (26 vs 43 known driver genes found) likely due to their introduction of increased noise of passenger mutations (**Supplementary Figure 5**). These data collectively show that ActiveDriverWGS accurately recovers known cancer driver genes and non-coding genome regions with frequent somatic mutations.

### Driver analysis reveals frequently mutated regulatory elements FMREs)

Having validated ActiveDriverWGS, we next sought to discover non-coding cancer drivers in *cis*-regulatory regions. We studied 4.5 million TFBS mapped in ENCODE^37^ in chromatin immunoprecipitation with DNA sequencing (ChIP-seq) experiments. We focused on 149,222 *cis*-regulatory modules (CRMs) that covered 103 Mbp and 3.3% of the genome. CRMs were defined by overlapping binding sites of at least two TFs that were observed in least two cell lines. To avoid confounding functional impact, CRMs segments overlapping coding regions and splice sites were excluded. The majority of CRMs (75%) overlapped with no UTR or promoter of protein-coding gene, enhancer or lncRNA sequence studied in *PCAWG^[PCAWG-2-5-9-14]^* (**Supplementary Figure 6**). These experimentally defined CRMs represent less-explored genomic space for driver discovery and are complementary to commonly used gene-focused regions such as fixed upstream promoters. The merging of overlapping TFBSs allowed us to reduce the redundancy of binding patterns of different TFs, while filtering of cell-type specific TFBSs led to a high-confidence set of regulatory regions more likely characteristic of a heterogeneous pan-cancer cohort of tumor samples.

Pan-cancer analysis of CRMs using ActiveDriverWGS revealed 41 frequently mutated regulatory elements (FMRE; *FDR*<0.05) (**Figure 1c, Supplementary Table 1**). FMREs included previously described recurrently mutated regions (promoters of *TERT* and *WDR74;* lncRNA *MALAT1*), serving as positive controls. Driver analyses of individual cancer types revealed six FMREs, including three not seen in pan-cancer results (**Supplementary Figure 7**). We found that FMREs were longer than CRMs in general (median 1049 bp vs 491 bp, Wilcoxon *P*=3.1×10^−6^) and included more TF binding sites (34 vs 10, *P*=2.1×10^−5^) while length and TFBS abundance were strongly correlated (Pearson r=0.64, *P*<10^−300^). The FMREs represented 698 patients (38%) with 1,092 SNVs and 113 indels. Most FMREs (25/41) did not overlap any UTRs, promoters, or lncRNA genes, including five intronic FMREs and 10 FMREs that were more than 50 kbp from any gene or annotated region. Thus our findings are complementary to gene-focused driver analyses in *PCAWG^[PCAWG-2-5-9-14]^*.

To confirm these findings, we used four additional methods MutSigCV^16^, NBR^41^, OncoDriveFML^42^ and DriverPower^[Shuai & Stein]^ that use distinct statistical models, clustering of mutations, and functional impact scores to find coding and non-coding cancer drivers. The majority of FMREs detected by ActiveDriverWGS (26/41) were also found by at least one other method, significantly more than expected from chance alone (0 expected, Fisher’s exact *P*=1.8×10^−77^). The five methods revealed a total of 92 candidate regions at *FDR*<0.05 and the FMRE at the *TERT* promoter was identified by all methods (**Figure 1d, Supplementary Figure 8**). Recovery of most FMREs with independent analytical approaches supports our findings of FMREs and suggests that some may act as cancer drivers that are subject to positive selection. However their elevated mutation frequency may also reflect regionalized hyper-mutation or challenging genomic regions with technical sequencing artefacts.

Power analysis suggests that FMREs with relatively rare mutations are only discoverable in large patient cohorts (**Supplementary Figure 9**). The PCAWG pancancer dataset is suitable for detecting effects three-folder smaller than for the largest PCAWG tumour-type specific cohorts (i.e. breast, prostate and pancreatic). We show that FMREs exist, but have been below the detectable effect-size in the larger individual tumor-type studies published to date. Thus we need to use pancancer analyses and sequence larger cancer-specific cohorts in the future.

### FMREs are enriched in multiple classes of somatic alterations

Cancer driver genes are affected by different genetic mechanisms in different tumors and tumor types. To further study the biological importance of FMREs, we analysed their somatic copy number alteration (CNA) and structural variation (SV) landscapes profiled in *PCAWG^[PCAWG-6; PCAWG-1]^* relative to expected genetic alterations. TF binding sites have been shown to have higher somatic mutation burden, potentially due to collisions of gene regulatory and DNA repair pathways. To account for TF occupancy as a cofactor of mutation rates, we sampled control regions from all CRMs according to their mean TF occupancy per nucleotide in 100 equally sized bins and used sampled CRMs to establish expected number of mutations according to the bin distribution of FMREs. As additional controls, we sampled regions with matching length randomly from the genome. To avoid biasing our analyses by earlier findings of recurrent non-coding cancer mutations and known drivers, we excluded 3 of 41 FMREs corresponding to the *TERT* promoter, the 5’UTR region of *WDR74* and the lncRNA *MALAT1*.

As a confirmation of ActiveDriverWGS analysis, FMREs as a group included significantly more SNVs and indels (880) than expected from all CRMs with similar TF binding occupancy (288 expected, *P_CRM_*=5.9×10^−5^) and from random genome-wide regions (113 expected; *P_GW_*<10^−6^) (**Figure 2a**). The enrichment suggests the mutations apparent in FMREs exceeds the mutation rate of comparable regulatory regions and may instead reflect positive selection of mutations important in cancer biology. Similarly, 96 structural variant breakpoints were significantly enriched in FMREs compared to both types of control regions (*P_CRM_*<10^−6^, 5 expected; *P_GW_*=6.0×10^−6^, 9 expected)(**Figure 2a**). Focal copy number variants (652) showed a trend of enrichment (*P_CRM_*=0.074, 511 expected; *P_GW_*=0.0081, 430 expected) (**Figure 2a**). In total, 43% of all patients in the dataset (793/1,844) had at least one mutation in any FMRE (SNV, indel, SV or focal CNA), significantly more than expected by chance from the distribution of TF-occupancy weighted CRMs (399/1,844; *P_CRM_*<10^−6^) or from random genomic control regions (469/1,844, *P_GW_*=1.1×10^−4^). Individual FMREs with fewer SNVs and indels often included many focal CNAs in additional patients, while few patients (46/793 or 6%) carried multiple mutations of different types in the same FMRE (**Figure 2b**). Thus FMREs likely include functionally important regions that are modified through distinct genetic mechanisms in different tumors and tumor types. For example, enrichment of structural variants among FMREs may indicate enhancer hijacking events mediated by translations^35,36^.

**Figure 2.**
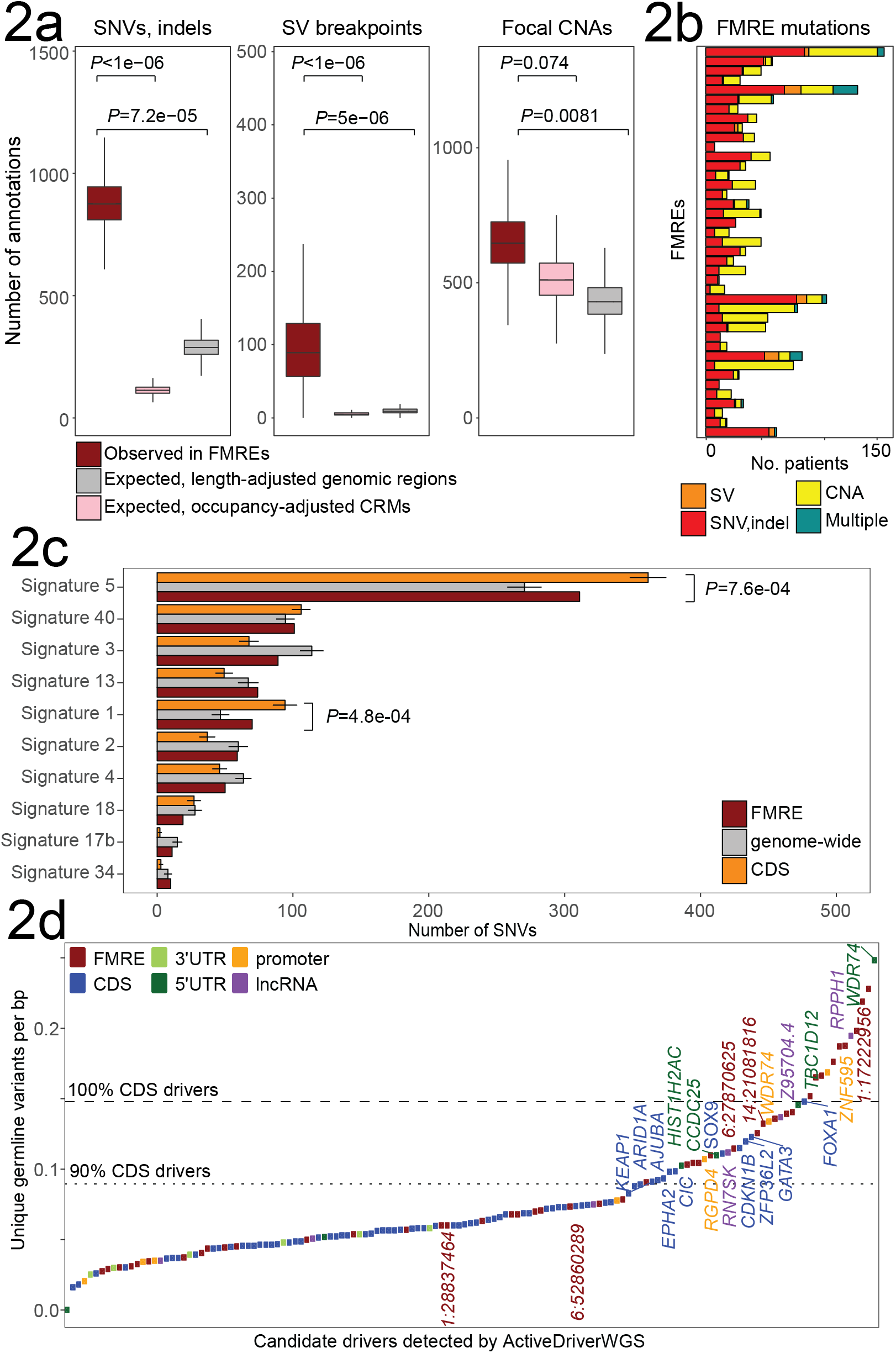
FMREs are enriched in different types of somatic mutations, aging-related mutation signatures and germline variants. (**a**) FMREs as a set are enriched in SNVs and indels, structural variation breakpoints and focal copy number alterations (dark red boxplots). As controls we used sets of CRMs sampled with matching average TF occupancy (pink boxplots) and sets of randomly sampled genomic regions (grey boxplots). Bootstrap resampling was used to estimate variation of FMRE mutations. FMREs corresponding to three previously known regions (*TERT, WDR74, MALAT1*) were excluded to estimate the properties of novel candidates and remove bias towards known regions. (**b**) FMREs involve distinct types of somatic alterations in different tumors and tumor types, while few FMREs carry multiple types of mutations in the same tumor. FMREs are ranked according to their significance in ActiveDriverWGS analysis. (**c**) Mutation signatures of SNVs in FMREs (dark red) are compared to signatures in protein-coding driver genes randomly sampled mutations. FMRE-associated mutations are enriched in aging-related signatures five and one, relative to randomly sampled mutations in the tumor samples with FMRE mutations. Error bars show one standard deviation above and below mean. (**d**) All regions identified by ActiveDriverWGS at *FDR*<0.05 ranked according to mean number of distinct SNVs per base pair. Genes with high germline variation and highlighted FMREs are labelled. Known protein-coding drivers detected by ActiveDriverWGS were used to estimate expected germline variation as 90^th^ and 100^th^ percentile (dashed and dotted line, respectively).

To study mutational processes active in FMREs, we evaluated the mutation signatures of SNVs using sample-specific exposure predictions developed by PCAWG^*[pcawg-7]*^ (**Figure 2c**). As controls, we sampled genome-wide mutations from the samples that carried FMRE mutations. We found that FMRE mutations were significantly enriched in aging-related signatures: signature five with 311 SNVs (permutation *P*=7.6×10^−4^, 270 SNVs expected) and signature one with 70 SNVs (*P*=4.8×10^−4^, 47 SNVs expected), relative to mutations sampled randomly from the tumor genomes with FMRE mutations. As expected, 59 known protein-coding drivers detected by ActiveDriverWGS were also enriched in signatures one and five relative to genome-wide mutations. The overall higher frequency of SNVs with aging-related signatures supports the hypothesis of FMREs acting as cancer drivers.

To reveal the FMREs with the strongest indications of hyper-mutation or technical biases, we studied germline variants in the PCAWG cohort (**Figure 2d**). FMREs had significantly more unique germline SNPs per nucleotide compared to exons of 59 protein-coding drivers (median 0.074 vs 0.058, Wilcoxon *P*=0.010), in agreement with recent findings of reduced mutation rates in exons due to differential mismatch repair^43^. Twelve frequently mutated regions, including nine FMREs (22%) as well as 5’UTR of *WDR74*, promoter of *ZNF595* and the lncRNA *RPPH1* exceeded the germline density of all protein-coding drivers and we flagged these as potetntially problematic (**Figure 1c**). Eleven additional FMREs (27%) lied between the 90^th^ and the 100^th^ percentile of germline variation of protein-coding drivers, similarly to known cancer genes (e.g. *FOXA1, GATA3*) and genes with cancer predisposition variants (e.g. *CDKN1B*). We also compared FMREs to common fragile sites^44^ and flagged five regions as potentially problematic, including two with excess germline variation in PCAWG. Thus driver discovery of non-coding regions such as CRMs is challenged by germline variation with biological and technical cofactors.

However some regions may also undergo positive selection in somatic genomes and include cancer predisposition variants in the germline genomes of cancer patients.

### FMREs are enriched in long-range chromatin interactions and superenhancers

To explore the potential role of FMREs as distal regulatory elements interacting with promoters of target genes, we studied chromatin long-range interactions representing the three-dimensional architecture of the genome. We annotated FMREs using loop anchors of 11,282 high-confidence chromatin interactions conserved in at least two cell lines derived from a public HiC dataset^24^. We found that 13/38 FMREs associated with distal genomic regions through 29 long-range chromatin interactions (**Figure 3a**). This is a two-fold enrichment relative to occupancy-matched CRMs (*P_CRM_*=0.0028, 13 interactions expected) and five-fold genome-wide enrichment (*P_GW_*=3.0×10^−6^, 6 interactions expected), suggesting that the mutated FMREs are particularly frequently interacting with distal genomic regions.

**Figure 3.**
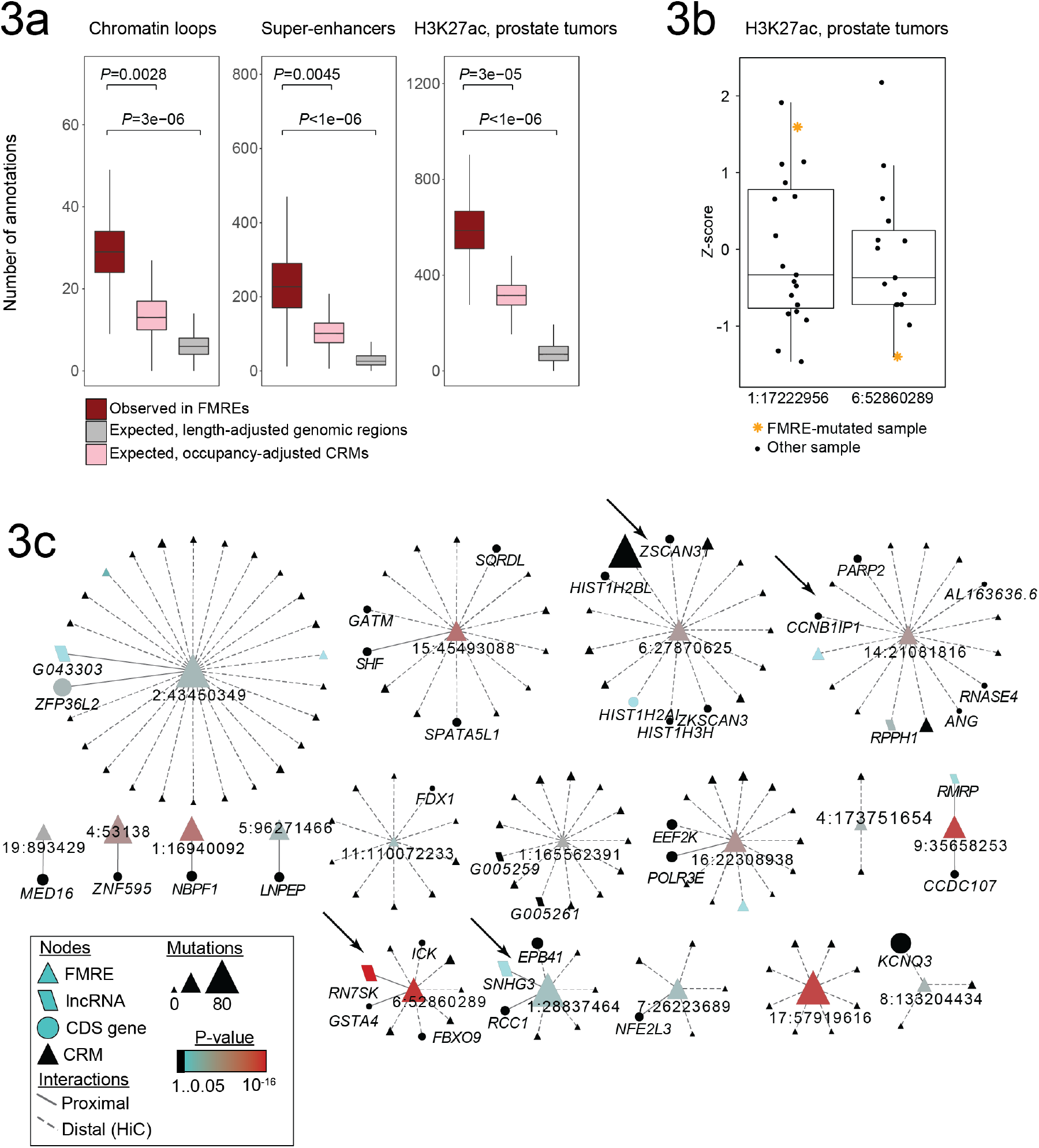
FMREs are enriched in super-enhancers and chromatin loops. (**a**) FMREs are enriched in long-range chromatin interactions of loop anchors, super-enhancer elements across multiple tissues, and enhancer histone marks (H3K27ac) of 19 primary prostate tumors with WGS data in PCAWG. Observed annotations in FMREs (dark red) are compared to TF occupancy-adjusted sampling of CRMs (pink) and genome-wide random regions (gray). (**b**) Two FMREs carry mutations that associate with stronger or weaker enhancer marks in primary prostate tumors. Boxplots show normalized H3K27ac signal in the FMRE near the mutation of interest. Yellow asterisks indicate the enhancer mark intensity in single samples with mutated FMREs. (**c**) Chromatin interaction network shows FMREs and their putative target genes. The network displays two types of interactions: proximal interactions comprise FMREs that coincide with gene promoters (solid line), and distal interactions comprise FMREs and genes that interact via chromatin loops (interactions) of promoters and FMREs (dashed line). Node size corresponds to number of mutations, color to mutation significance and shape to type of genomic region. Regions highlighted in the text are indicated with arrows.

To explore the potential role of FMREs as *cis*-regulatory elements, we used a dataset of 58,283 super-enhancers^45^ across 86 human cell types. Super-enhancers are sets of adjacent enhancers (also known as clusters of open regulatory elements (COREs)) that are bound by master regulators and involved in cell type specification^46,47^. Half of FMREs (19/38) occurred at 234 super-enhancers of various tissues and were enriched relative to both sets of control regions (*P_CRM_*<0.0045, 101 annotations expected; *P_GW_*<10^−6^, 26 expected) (**Figure 3b**). Tissue-specific superenhancers co-occurred with FMREs more frequently than expected with 31 tissue types (*P_CRM_*<0.1) including fetal cells, hematopoietic and immune cells, as well as five cancer cell lines (**Supplementary Figure 10**). In total, 25/38 FMREs were annotated at either super-enhancers or chromatin loop anchors and seven FMREs with both types of genomic elements, suggesting that mutations in FMREs rewire the cis-regulatory logic encoded by super-enhancers and their long-range chromatin interactions.

To validate our observations of enriched super-enhancers in FMREs, we studied a genome-wide ChIP-seq dataset of histone H3 lysine 27 acetylation (H3K27ac) sites representing active enhancers of 19 primary prostate cancer samples^48^ with matched WGS data in PCAWG. FMREs were significantly enriched in 591 H3K27ac peaks (*P_CRM_*<4.4×10^−5^, 315 sites expected; *P_GW_*<10^−6^, 69 sites expected) (**Figure 3c**). A sizeable portion of FMREs (18/38) appeared as active enhancers in the majority of prostate samples and most FMREs (25/38) showed ewnhancer marks in at least one prostate tumor sample of the subset (**Supplementary Figure 10**). These data support the hypothesis that mutations in FMREs are engaged in gene regulation in primary tumors.

We asked whether the mutations in FMREs associated with differential H3K27ac signal in the 19 H3K27ac-profiled prostate tumors. Of the five FMREs with mutations in relevant samples, two FMREs showed mutation-associated differences in H3K27ac levels. A single mutation in the FMRE 1:17222956 corresponded to the sample with the highest H3K27ac peak in the region (z-score=1.67; **Figure 3b**), while a mutation in the FMRE 6:52860289 corresponded to the sample with the lowest H3K27ac peak (z-score=-1.68; **Figure 3b**). Both FMREs were detected as candidate drivers by four driver discovery methods, while the first region was flagged due to excess germline variation. Although limited in statistical significance due to single mutated samples, these observations suggest that FMRE mutations may co-occur with altered chromatin marks.

Enrichment of FMREs in regions with chromatin interactions and superenhancer annotations suggests that FMREs and corresponding somatic mutations are involved in central gene regulatory programs of tissue identity and differentiation. Known and unknown regional mutational processes active in gene regulatory processes may confound our observations of candidate drivers, however the occupancy-weighted permutation procedure shows that FMREs are enriched in regulatory annotations beyond what is expected from other frequently TF-bound regions. Further analyses and experimental work is required to deconvolute the effects of somatic mutation rates and positive selection apparent in super-enhancers and chromatin interaction sites.

### Chromatin interactions of FMREs reveal mutation impact on gene expression

To study the impact of candidate driver mutations in FMREs, we associated FMREs and putative target genes using high-confidence chromatin interactions. The resulting chromatin interaction network included 18/38 FMREs and 37 putative target genes that either shared promoter or 5’UTR sequence with FMREs (15 genes) or were distally associated to FMREs via long-range chromosomal interactions (22 genes) (**Figure 3c**). The remaining 20 FMREs with no apparent target genes were excluded.

We tested associations of 11 FMREs and 22 potential target genes for differential gene expression and revealed seven (32%) genes (*RCC1, CCNB1IP1, GSTA4, ICK, HIST1H2AI, ANG, ZKSCAN3*) with differential mRNA abundance in samples with mutations in four FMREs (Chi-square *P*<0.05, *FDR*<0.14). We used the PCAWG transcription dataset^*[PCAWG-3]*^ that covered ~50% of samples with WGS data and applied negative binomial regression models on mRNA abundance values (RPKM-UQ) that controlled for cancer type and relative gene copy number variation as covariates. To increase confidence, we analyzed tumor types with at least three mutated samples and excluded genes with low mRNA abundance (mean RPKM-UQ > 1).

*CCNB1IP1*, a tumor suppressor gene according to the Cancer Gene Census database, showed reduced expression in three kidney and three breast tumors with available gene expression data (P=0.0083, **Figure 4a**). The FMRE 14:21081816 located 280kbp downstream of *CCNB1IP1* was mutated in 24 tumors in total (six expected by chance, *FDR*=6.2×10^−3^). The FMRE was detected as significant by three driver discovery methods. The 1.3 kbp FMRE interacts distally with *CCNB1IP1* through long-range chromatin interactions and is bound by 87 TFs in ENCODE, likely representing a high-occupancy target (HOT) region bound by dozens of TFs and involved in developmental enhancer function^49,50^. *CCNB1IP1* (cyclin B1 interacting protein 1) encodes a ubiquitin E3 ligase that negatively regulates cell motility and invasion by inhibiting cyclin B1^51,52^. The angiogenesis-related gene *ANG* interacting with the FMRE via chromatin loops also showed lower expression in FMRE-mutated samples (*P*=0.042).

**Figure 4.**
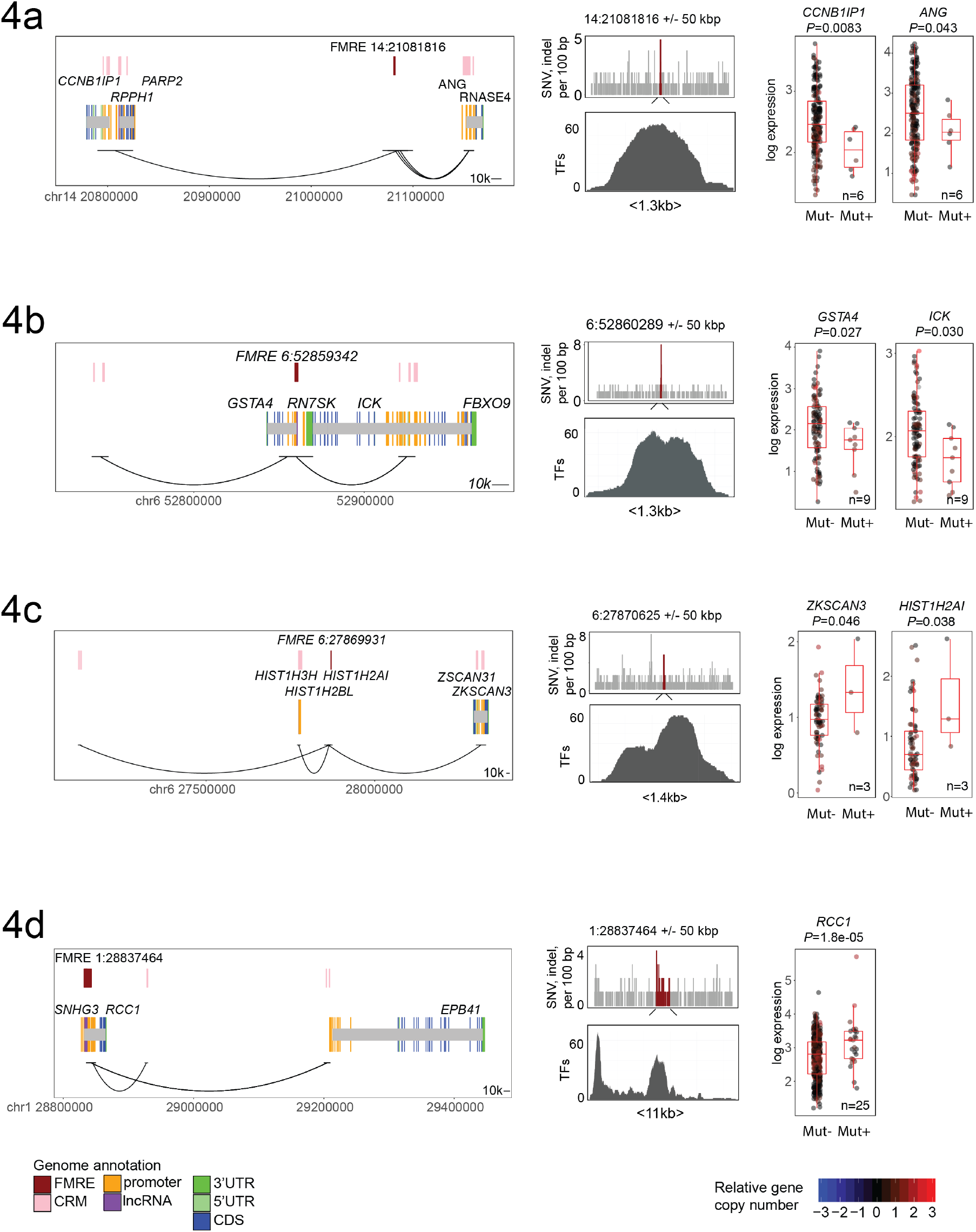
Mutations in FMREs associate with differential expression of cancer genes. Left: chromosomal location of FMREs, target genes and long-range chromatin interactions of gene promoters and FMREs. Middle: mutations in the FMRE and 50 kbp flanking region (top) and histogram of TF binding in the region (bottom). Right: altered expression of target genes in FMRE-mutated samples. Points represent log1p-transformed expression values (RPKM-UQ) and are colored according to relative copy number of target gene. (**a**) The tumor suppressor gene *CCNB1IP1* and angiogenesis related gene *ANG* showed reduced expression in six kidney and breast cancer samples with mutations in distal FMRE. (**b**) The drug metabolism gene *GSTA4* and intestinal kinase gene *ICK* showed reduced expression in nine breast and bladded cancer samples with mutations in the distal FMRE. (**c**) The candidate oncogenic transcription factor *ZKSCAN3* and histone gene *HIST1H2A1* showed increased expression in three ovarian cancer samples with mutations in the distal FMRE. (**c**) The novel cancer gene *RCC1* involved in RanGTP signaling and cell cycle shows increased expression in 25 samples of seven cancer types with mutations in the proximal FMRE upstream of the gene.

The genes *GSTA4* and *ICK* showed reduced expression in 6 breast and 3 bladder cancer samples with mutations in the FMRE 6:52860289 (*P*=0.027 and *P*=0.030 respectively) (**Figure 4b**). The FMRE has mutations in 33 samples (7 expected by chance; *FDR*=5.8×10^−9^), overlaps with the promoter of *GSTA4*, the small nuclear RNA *RN7SK*, and has long-range chromatin interactions with the promoter of *ICK. GSTA4* encodes the metabolic enzyme glutathione S-transferase alpha 4 involved in cellular defense against toxic, carcinogenic, and pharmacologic compounds and stress-induced TP53 signaling for apoptosis^53^. *ICK* encodes the intestinal cell kinase involved in cell cycle^54^ and implicated in proliferation and ciliogenesis in glioblastoma^55^. The FMRE is annotated as a super-enhancer in brain hippocampus and carries binding sites of 103 TFs. We first found this FMRE due to a mutation that associated with decreased H3K27ac level in prostate tumors (**Figure 3b**). Reduced expression of *GSTA4* and *ICK* and decreased level of the enhancer-associated histone mark in mutated samples fit the hypothesis that mutations at this FMRE disrupt gene expression.

The transcription factor *ZKSCAN3* showed increased abundance in three ovarian cancer samples with available gene expression data (chi-square *P*=0.046, **Figure 4c**). The FMRE 6:27870625 bp was mutated in 27 pan-cancer samples (8 expected, *FDR*=8.0×10^−4^) and was considered significant by two driver discovery methods. The 1.4 kbp region interacts with target genes through long-range chromatin interactions and includes a thymus-related super-enhancer and a HOT region bound by 74 TFs. *ZKSCAN3* (zinc finger with KRAB and SCAN domains 3) located ~450 kbp downstream of the FMRE is a transcriptional repressor of autophagy^56^ and a positive regulator of the cyclin D2 oncogene in multiple myeloma^57^. It has also been implicated in the promotion, migration and metastasis of colorectal^58,59^, prostate^60^, and bladder cancer^61^. The adjacent histone gene *HIST1H2AI* interacting with the FMRE via chromatin loops also showed differential expression relative to mutations in this FMRE (*P*=0.038).

The strongest association of FMRE mutations and mRNA abundance was found at the FMRE upstream of *RCC1* (regulator of chromosome condensation 1). *RCC1* showed elevated expression in 25 FMRE-mutated samples of bladder, breast, colorectal, kidney, lung and ovarian cancers (chi-square *P*=1.8×10^−5^, **Figure 4d**). The FMRE was mutated in 59 tumors in total (33 expected, FDR=0.0499). The FMRE 1:28837464 is a 11 kbp region that includes the *RCC1* promoter, the adjacent ncRNA *SNHG3*, binding sites of 102 TFs, and super-enhancers of cancer cell lines (liver HepG2; leukemia K652; colon HCT116) and hematopoietic and immune cells (CD4+, CD8+, CD34+). *RCC1* is not characterized in cancer, however its involvement in hallmark cancer pathways suggests it as a candidate oncogene. *RCC1* encodes a DNA-binding guanine nucleotide exchange factor that produces the RanGTP signaling molecule essential for mitotic processes^62–64^. *RCC1* is regulated by MYC^65^ and its overexpression in normal cells evades DNA damage-induced cell cycle arrest and senescence^66^.

To increase confidence in these candidate drivers, we manually reviewed all 148 mutations in raw sequencing data files and evaluated their sequence context, read coverage and strand bias. The majority of all mutations (142 or 96%) and all mutations with matching expression data (56) were considered true positives while 17% of mutations (25/142 and 10/56) were flagged due to strand bias or low variant allele frequency. The false positive rate corresponds to overall variant calling error rate of the PCAWG project.

In summary, these examples suggest that a subset of non-coding mutations in FMREs increase oncogenic gene expression or reduce the transcription of tumor suppressor genes, further supporting their role as candidate cancer drivers. Alternative definitions of tissue-specific regulatory elements, gene regulatory regions and chromatin interactions detected in primary tumor samples of matching tissue types, and larger datasets of matched transciptomes will likely reveal further FMREs and target genes.

### FMRE mutations at *RCC1* locus associate with global activation of immune response genes

The large number of mutations in the FMRE upstream of *RCC1* prompted us to study global differential gene expression of mutated and non-mutated cancer samples across 9,420 protein-coding genes with Gene Ontology (GO) annotations and above-baseline transcript abundance using the cancer type and copy number adjusted statistical models described above.

We found 62 significantly expressed genes (*FDR*<0.05, chi-square test), all of which showed increased expression in FMRE-mutated samples relative to nonmutated samples of matched cancer types (**Figure 5a**). To further characterize the genes up-regulated in FMRE-mutated tumors, we carried out pathway enrichment analysis of FDR-ranked genes using g:Profiler^67^ and found 16 biological processes of GO and 3 Reactome pathways (*FDR*<0.05). Intriguingly, 34/62 differentially expressed genes were significantly enriched in immune response, neoantigen processing, endocytosis and fiber elongation pathways (**Figure 5b**). The activation of immune response genes and pathways such as *antigen processing and presentation (WAS, SLC11A1, CAPZB, LILRB2, RFTN1, CTSL, CCL19, AP1S2; FDR*=0.0024) is in agreement with the super-enhancer annotations of hematopoietic and immune cells associated with the FMRE. The differentially genes also included one known cancer gene *WAS* implicated in the Wiskott-Aldrich immunodeficiency syndrome that has been associated with lymphoma^68^. While further experimental work is required to elucidate the underlying mechanisms, our differential expression and pathway analysis suggests that the cancer mutations in the FMRE upstream of *RCC1* activate global gene expression patterns, potentially to enhance the activity of hallmark cancer pathways of immune suppression.

**Figure 5.**
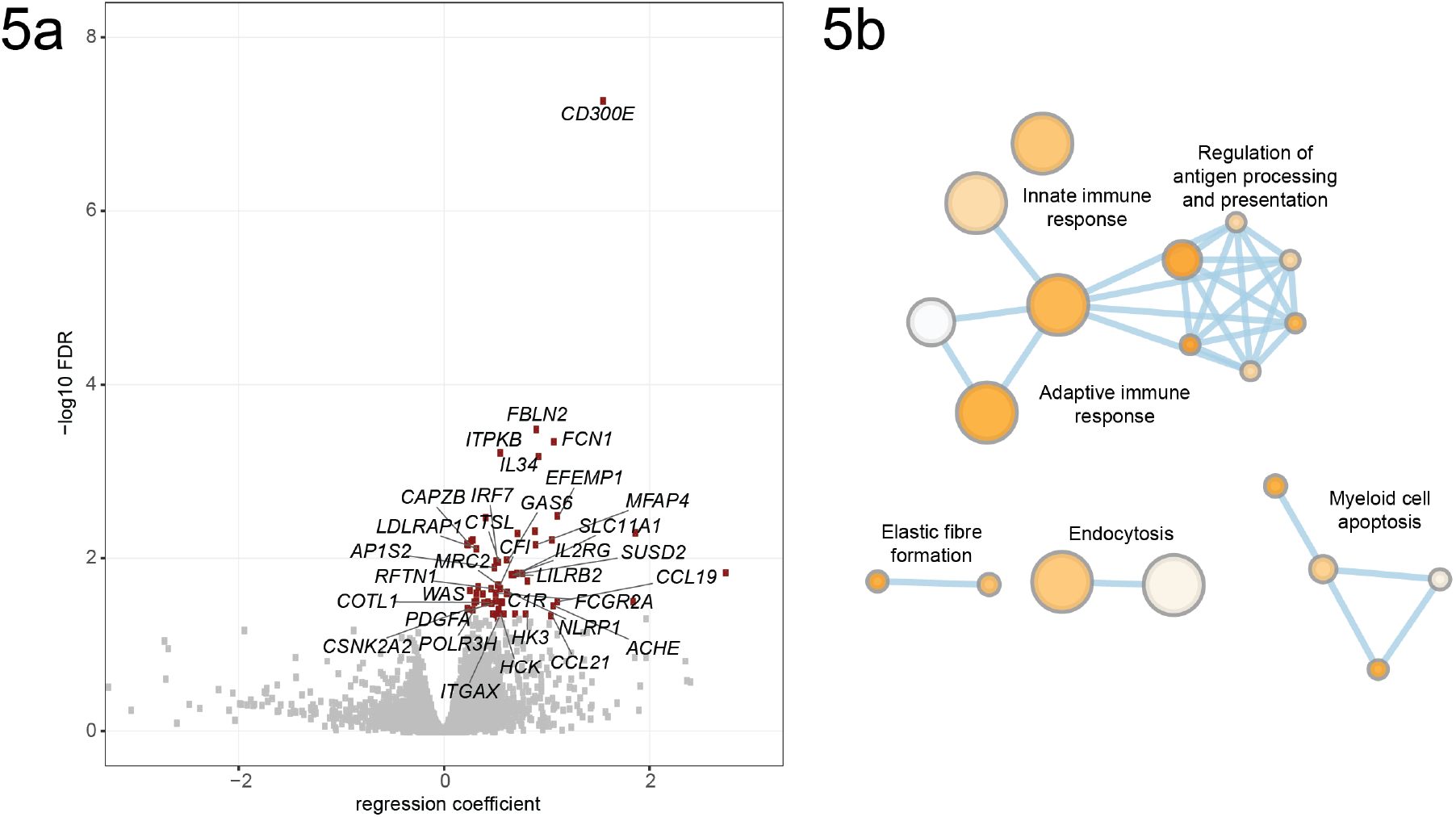
Mutations in FMREs at *RCC1* locus associate with global activation of immune response pathways. (**a**) Volcano plot shows genes with differential expression in tumors with mutations in the FMRE upstream of the *RCC1* gene. Genes with significant expression differences are shown in dark red (*FDR*<0.05) and gene symbols with enriched pathway annotations are shown. (**b**) Enrichment map shows significantly enriched GO processes and Reactome pathways corresponding to enriched genes (*FDR*<0.05 from g:Profiler). Network nodes represent pathways and processes and nodes with many shared genes are connected with edges. Subnetworks are annotated with common biological themes representative of pathways.

## Discussion

Only few non-coding cancer drivers are known to date. Their discovery requires large WGS datasets and detailed annotations of the regulatory genome. Thus the search space of driver discovery efforts has been limited to gene-focused regions of the genome. Here we performed a driver analysis of the *cis*-regulatory genome using the largest cancer WGS dataset available to date from the PCAWG project. We revealed the currently largest set of pan-cancer driver candidates, frequently mutated regulatory elements (FMREs), that were enriched in somatic non-coding SNVs and other genomic alterations across a heterogeneous cohort of tumors. Two thirds of FMREs occurred at known super-enhancers or chromatin loops and most appeared as enhancers in primary tumors. Our leading hypothesis is the positive selection of these regions in cancer genomes that causes oncogenic rewiring of gene regulatory networks and long-range chromatin interactions of distal enhancers and target genes. We found several lines of evidence support the driver hypothesis: enrichment of different classes of mutations in FMREs, over-representation of aging-associated mutation signatures, and significant associations of candidate driver mutations and expression of putative target genes and pathways involved in hallmark cancer processes.

We cannot rule out alternative explanations to observed enrichment of somatic mutations in the identified regions. Thus caution should be taken in interpreting these candidate driver regions, most of which are reported for the first time. From the point of genome biology, the somatic mutation landscape has complex associations with chromatin state and gene regulation. While open chromatin is broadly associated with reduced mutation load, abundant mutations in TF-bound regions have been associated with deficient DNA repair due to competitive binding of regulatory and DNA repair proteins. However our analysis shows that FMREs are enriched in mutations and regulatory annotations even when considering regions with similar TF occupancy as controls, suggesting that the observed mutation enrichment may be due to positive selection. Technically, the non-coding genome includes challenging regions with potential for sequence alignment and variant calling artefacts. We rely on the comprehensive preprocessing and filtering pipeline of the PCAWG project that uses a consensus of several state-of-the-art methods for variant calling. Some FMREs have high germline variation that potentially originates from highly variable regions such as fragile sites, regions that are challenging the sequencing pipeline, as well as regions with functional germline variants of cancer predisposition. Further computational analyses and experimental work are required to establish these candidate noncoding regions as *bona fide* cancer drivers.

To capture and interpret pan-cancer drivers, we analysed high-confidence regulatory regions and long-range chromatin interactions apparent across multiple cell lines. These regions and interactions are more likely representative of a pancancer cohort than those of single cell lines, however any epigenomic data derived from cell lines are limited in their biological relevance to primary tumors. Thus future driver analyses of non-coding regions will benefit from epigenomic and gene regulatory profiles derived from matching tumors and tumor types.

Our analysis revealed rarely mutated FMREs that were detectable only in the pan-cancer dataset while few cancer type specific FMREs beyond the *TERT* promoter were identified. Our power analysis confirms that the available sample sizes do not permit analysis within cancer types and suggests that considerably larger tumor cohorts with WGS data are required for future studies. Discovering functional driver mutations in FMREs using target gene expression was even more limited as only half of PCAWG tumors had matching transcriptomic data available. Thus additional FMREs likely remain to be discovered.

Integration of cancer genome variation with epigenomic profiles, long-range chromatin interactions and matching transcriptomic data is a powerful approach for discovering candidate drivers and mechanistic hypotheses of the roles of mutations. This strategy is applicable to tissue-specific regulatory regions as well as other types of regions such as ultra-conserved elements. Systematic genetic disruption of candidate driver regions with the CRISPR technology coupled with phenotypic screens is required to demonstrate the function of mutations in FMREs in cell lines and model organisms. Analysis of future WGS datasets paired with comprehensive clinical information such as those generated in the ICGC-ARGO project will enable biomarker discovery from non-coding mutations. In summary, our study suggests that the non-coding cancer genome includes previously uncharacterized rare driver mutations that contribute to the hallmarks of cancer through cis-regulatory mechanisms. Further computational and experimental studies are needed to understand the role of these regions and the non-coding cancer genome with its mutational processes and driver mechanisms.

## Methods

### Somatic mutations

We analyzed the dataset of 1,844 whole cancer genomes of 31 cancer types with 14.2 million somatic single nucleotide variants (SNVs) and *indels^[PCAWG-1]^*. This represented a subset of the consensus dataset of 46.6 million mutations in 2,583 samples sequenced in the Pan-cancer Analysis of Whole Genomes (PCAWG) project of the International Cancer Genome Consortium (ICGC). The subset was derived using the following procedure. First we filtered 69 hyper-mutated samples with more than 90,000 mutations (~30 mutations/Mb) that contributed 47% of all mutations. We further excluded 670 samples of four cancer types: melanoma (65), lymph-related cancers (BNHL (104), CLL (90), NOS (2)), esophageal adenocarcinoma (95), and liver hepatocellular carcinoma (314) to avoid leakage of stronger mutation enrichment signal of these cancer types to the pan-cancer cohort (see **Supplementary Note 1**).

### Genomic regions

Our driver discovery pipeline was run separately for multiple classes of genomic regions of the human genome hg19. Cis-regulatory modules from the ENCODE project comprised clusters of transcription factor (TF) binding sites (TFBS) measured in chromatin immunoprecipitation (ChIP-seq) experiments retrieved from UCSC Genome Browser. We used the dataset of 4.9 million binding sites of 161 TFs in 91 cell lines and excluded sites that were only observed in one cell line. The remaining 1.1 million binding sites of 101 TFs were merged into consecutive regions based on ≥1bp of common sequence, resulting in 322,614 regions. We discarded regions bound by single TFs and used the remaining 149,222 clusters of TFBS (*i.e*., cis-regulatory modules, CRMs) for driver discovery. CRMs were filtered to exclude sequence regions overlapping with coding sequence and splice sites. In addition to CRMs, we performed driver discovery on protein-coding sequences (CDS), untranslated regions of protein-coding genes (5’UTR, 3’UTR), promoters of coding genes (promDomain), and gene bodies of long non-coding RNAs (lncRNA) derived from the PCAWG consensus *dataset^[PCAWG-2-5-9-14]^*.

### ActiveDriverWGS and driver discovery

Candidate cancer driver genes and regulatory regions were identified with ActiveDriverWGS, our novel mutation enrichment method that tests whether a genomic element of interest is significantly more mutated than the relevant background sequence using a generalized linear regression model. ActiveDriverWGS is a local mutation enrichment model that determines the expected number of mutations in a genomic region by observing mutations in a background window of at least 100kb around the region of interest, including ±50kb upstream and downstream of the region plus additional intermediate regions such as gene introns. ActiveDriverWGS considers sequence trinucleotide composition as a cofactor in the regression model. It models the number of all sequence positions of each of 32 classes of trinucleotides in both the background sequence and sequence region of interest as well as the number of mutations in these trinucleotide classes. Indel mutations are modeled as the 33^rd^ class of mutations with equal probability at each sequence location. Only one mutation is counted per tumor in cases where an element contains multiple mutations in the same tumor genome. This reduces the impact of local hypermutations and leads to more conservative driver prediction. ActiveDriverWGS conducts chi-square tests to validate two hypotheses using pairs of hierarchical regression models (*H0 vs. H1*). The statistical test checks whether mutations in the region of interest (variable *is_element*) are distributed differently relative to its background sequence:

*H0:* n_mutations ~ Pois (trinucleotide_context)
*H1:* n_mutations ~ Pois (trinucleotide_context + is_element)

A significant p-value in this combined test indicated that the element of interest was a candidate cancer driver. To distinguish regions with excess mutations from regions with fewer than expected mutations, we additionally computed confidence intervals to expected numbers of mutations from the null model *H0* and accepted the alternative hypothesis *H1* only if the expected background mutations were significantly fewer than observed mutations at 95% quantile. If the confidence intervals indicated significant excess of mutations in the background and depletion in the region of interest, we inverted corresponding small p-values (*P*=1-*P*). Regions with no mutations were assigned *P*=1. The p-values resulting from the first test were corrected for multiple testing across all tested regions using the Benjamini-Hochberg False Discovery Rate (FDR) procedure and genes with *FDR*<0.05 were considered significant. The p-values from the second test were also corrected with the FDR procedure, limiting to elements that passed the first test at *FDR*<0.05. Each cancer type and element type was subject to separate multiple testing correction procedure.

Power calculations for chi-squared tests in ActiveDriverWGS were conducted using the pwr.chisq.test function of the ‘pwr’ package in R. Effect size was computed using number of samples, final degrees of freedom from ActiveDriverWGS output (1), and significance level (*P*=0.05). This process was repeated for several values of power (0.6–0.9) and data were plotted as line plots.

The R source code of ActiveDriverWGS is freely available at https://github.com/reimandlab/ActiveDriverWGS.

### Benchmarking of ActiveDriverWGS

We tested ActiveDriverWGS using simulated mutations and parameter settings. To generate simulated mutation data, we split the genome into 50kb windows and randomly re-assigned PCAWG pan-cancer single nucleotide variants in each window to alternative positions of the same trinucleotide context using sampling with replacement. Indels were randomly re-assigned without using trinucleotide context. Besides in-house simulated data, we also tested ActiveDriverWGS on two additional sets of simulations from the PCAWG drivers group (Sanger, Broad). In total 672 simulation runs with three sets of simulated mutations, 32 cancer types and seven types of genomic elements revealed eleven significant findings at FDR<0.05, suggesting that very little deviation existed from expected false discovery rates. We also tested ActiveDriverWGS with different sizes of background windows: ±10kb, ±25kb, ±50kb, ±75kb, and ±100kb. We found that the method is robust to variations in background window, however the ±50kb window provided the best accuracy and enrichment of known cancer genes. We also excluded the trinucleotide cofactor in our regression models and observed a large increase in false positive findings. We repeated the analysis after including hyper-mutated samples and found that many fewer known driver genes were detected. Thus hyper-mutated samples were excluded from the analysis.

### Additional driver discovery methods

Four independent driver discovery methods were used to discover candidate drivers among CRMs. Each method used different statistical models, cofactors, mutation impact scores and/or clustering metrics to find candidate drivers. ***NBR*** uses a negative binomial regression to estimate the background mutation rate of each element as described earlier^41^. This method accounts for the length of each element and its mutability using a trinucleotide substitution model with 192 rate parameters and uses the local mutation rate in regions around each element as a covariate. ***DriverPower*** DriverPower is a combined burden and functional impact test for coding and non-coding cancer driver elements. In the DriverPower framework, randomized non-coding genome elements are used as training set. In total 1373 reference features covering nucleotide compositions, conservation, replication timing, expression levels, epigenomic marks and compartments are collected from public databases for downstream modelling. For the modelling, a feature selection step by randomized Lasso is performed at first. Then the expected background mutation rate is estimated with selected highly important features by binomial generalized linear model. The predicted mutation rate is further calibrated with functional impact scores measured by LINSIGHT^69^ scores. Finally, a p-value is generated for each test element by binomial test with the alternative hypothesis that the observed mutation rate is higher than the adjusted mutation rate, and the Benjamini-Hochberg procedure is used for FDR control. ***OncoDriveFML***, Driver discovery with OncoDriveFML was performed as described in the PCAWG driver study ^*[pCAWG-2-5-9-14]*^. ***MutSigCV***. Driver discovery with MutSigCV was performed as described in the PCAWG driver study ^*[pCAWG-2-5-9-14]*^.

### Super-enhancers and long-range chromatin interactions

We annotated FMREs using public datasets of long-range chromatin interactions and super-enhancers. The super-enhancer dataset originates from the study by Hnisz *et al*^45^. Chromatin loops representing long-range interactions from eight human cell lines were derived from the HiC dataset by Rao *et al*^24^. To obtain a high-confidence set of chromatin interactions, we merged interactions whose loop anchors overlapped with each other at both ends, and filtered those interactions that had been characterized only in one cell line. Long-range chromatin interactions were considered to interact with a gene if one anchor of the loop overlapped the coding, UTR or promoter sequence of the gene while the other anchor of the loop had no overlap with the gene. We also tested the aggregated set of H3K27ac and DNAse sites from the Roadmap Epigenomics project^70^. To determine statistical significance of genomic annotations of FMREs, we tested the union of all sequences corresponding to anchors using the two permutation strategies described below.

### Enrichment of regulatory annotations of FMREs

We counted the number of pairs of FMREs and distinct genomic annotations. To determine the statistical significance of enriched genomic annotations of FMREs, we used a custom permutation test to sample from all CRMs from ENCODE as controls. We split our initial dataset of ~150,000 CRMs into 100 equal bins based on their TF occupancy, represented as number of TFs bound in CRM divided by length of region. To estimate the expected number of regulatory annotations in FMREs, we sampled 10,000,000 random sets of CRMs from the bins using the number and size distribution of detected FMREs. Statistical significance of enriched annotations was estimated as an empirical p-value, i.e., the fraction of 10,000,000 permutations that showed equivalent or higher number of regulatory annotations than associated with the true set of FMREs. To avoid biasing our findings by known non-coding drivers, we excluded three FMREs overlapping with the *TERT* promoter, the *WDR74* promoter and the lncRNA *MALAT1*. Besides length-adjusted sampling of CRMs, we also sampled random genome regions of equivalent sizes as controls. Confidence intervals for observed numbers of FMRE annotations were derived with resampling.

### Copy number alterations and structural variants

Matching copy number and structural variation datasets originate from the PCAWG project^*[PCAWG-6;PCAWG-11]*^. We determined relative digital copy numbers of all regions and patients by accounting for previously computed sample ploidy estimates, whole genome duplication events, and patient sex. To estimate the frequency and enrichment of copy number alteration events in FMREs, we focused on focal and potentially high-impact copy number alterations with less than 5 mbp in size and total copy number of zero (corresponding to homozygous deletion) or relative gain of two or more copies. For structural variants, we studied coordinates of breakpoints. To determine statistical significance, we used permutation tests relative to all occupancy-matched ENCODE CRMs as well as size-matched random regions from the genome, using the strategy defined above. For analysis of mutation impact on gene expression, copy number altered regions were further processed to obtain gene-level copy number estimates. Copy numbers of genes were computed as the most extreme copy number values of their exons.

### Mutation signature analysis

To analyse mutation signatures characteristic of FMREs, we studied sample-specific exposure predictions for SNVs developed by PCAWG^*[PCAWG-7]*^. As controls, we sampled two sets of mutations in the cancer samples with FMRE mutations: SNVs present in 59 protein-coding drivers predicted by ActiveDriverWGS, and genome-wide SNVs. We conducted custom permutation tests with 100,000 sets of SNVs that were sampled with replacement using the number of mutations observed in FMREs and their distribution among cancer types. We computed the enrichment of each FMRE-related mutation signature by counting the fraction of randomly sampled genome-wide sets of SNVs that exceeded the number of SNVs observed in FMREs. Empirical enrichment p-values were derived as the fraction of permutations where sampled mutations of signature-specific SNVs exceeded the number of observed signature-specific SNVs in FMREs. Mutation signatures with fewer than 5% of genome-wide permutations exceeding observed FMRE signatures were highlighted as enriched.

### Germline analysis

Germline variant frequency of FMREs was estimated using density of unique SNVs and indels in 100 bp windows across the entire PCAWG cohort of cancer patients. As reference we used the protein-coding drivers identified by ActiveDriverWGS in the pan-cancer cohort. We computed germline variant density within genomic elements and estimated the upper bound of expected variation for within-element variation as 90^th^ and 100^th^ percentiles of values observed among coding driver predictions. FMREs that exceeded the 100^th^ percentile threshold were flagged for excess germline variation. We also computed germline variation density for frequently mutated promoters, UTRs, enhancers and lncRNAs discovered by ActiveDriverWGS in the pan-cancer driver analysis.

### Mutation impact on gene expression

We used matching RNAseq gene expression data from PCAWG^*[PCAWG-3; PCAWG-14]*^ to estimate the impact of non-coding mutations on gene expression. We used upper-quartile normalization values of fragments per kilobase of transcript per million (FPKM-UQ) as gene expression measurements. Approximately 50% of PCAWG tumors with WGS data had corresponding transcriptomic data, and the tumors with no available transcriptomic data were excluded from the analysis. Negative binomial regression models similar to ActiveDriverWGS were used to determine whether mutations in FMREs corresponded to significantly lower or higher gene expression levels in matching samples. The model evaluated gene rounded RPKM-UQ values and accounted for cancer type and relative gene copy number as cofactors. The alternative model tested whether mutated samples showed significantly different gene expression of gene of interest. In each statistical test, only samples with matching mutation, gene expression and copy number data were included and other samples were excluded. Further, cancer types with at least three mutated samples were included and others were excluded. Cancer types with fewer mutated samples were also removed from the control (non-mutated) set. Each FMRE was tested using pan-cancer mutations with cancer type as a nominal co-factor and relative copy number as a numeric cofactor. Two sets of genes were tested for every FMRE: genes or their promoters directly overlapping with an FMRE, and genes distally associated with an FMRE via a long-range chromatin interaction of gene promoter. Genes with *P*<0.05 were selected as significant and FDR values were computed across all tested pairs and reported (*FDR*<0.15).

### Global gene expression and pathway enrichment analysis

Global analysis of gene expression in samples with mutations in the FMRE upstream of *RCC1* was conducted with the same statistical approach as for single target genes. We tested protein-coding genes that had at least one GO annotation and showed above-baseline gene expression in tested samples (mean RPKM-UQ transcript abundance values above one unit). Genes with *FDR*<0.05 were selected as significant. Pathway enrichment analysis of genes ranked by p-values was carried out using the g:Profiler^67^ R package with the following settings: ranked input gene list, only GO biological processes and Reactome pathways considered, minimum five and maximum 1000 genes per gene set, g:Profiler internal multiple testing correction used for FDR estimates, minimum three genes shared with gene list and gene set, and electronic gene annotations (IEA) included.

### Mutation vetting

Mini-BAM files for samples with a variant in the following FMREs were downloaded from GNOS: chr1:28831933-28842995, chr6:52859342-52861236, chr6:27869931-27871319 and chr14:21081147-21082486. FMRE variants were manually examined in the IGV software v2.3.97. Variants that were missing or were called within palindromic regions were marked as false positives. Variants were flagged as low confidence if they occurred on one strand (forward or reverse), had fewer than four reads, or were found within a homopolymer run. Variants were highlighted, but not flagged, if they had four supporting reads, only one supporting read on one of the strands, in a low coverage region, or in a region with strand bias. Variants found in regions with strand bias, and showed strand bias in their supporting reads were highlighted, but not flagged.

### ChlP-seq data of primary prostate cancers

We used a recent ChiP-seq dataset for the histone mark H3K27ac in 19 PCAWG prostate cancer samples^48^. We examined the dataset to find overlaps of FMREs and H3K27ac peaks. Global overlap of FMREs and H3K27ac peaks was determined using the permutation strategy and the two types of control regions described above. To evaluate mutation impact on specific H3K27ac peaks within FMREs, the FMREs were extended by 1Kb up and downstream, and rounded to the nearest 100 bp before intersecting with H3K27ac regions determined in ChiP-seq files. Patients with an H3K27ac peak in the target region were considered to have an enhancer mark in proximity to the FMRE. Peak scores were subsequently converted to z-scores and plotted as boxplots.

## Acknowledgements

We would like to thank Federico Abascal, Gary Bader, Peter Campbell, Gad Getz, Nuria Lopez-Bigas, Inigo Martincorena, Jakob Skou Pedersen, Esther Rheinbay, and Josh Stuart for constructive comments on the project and the manuscript, Julian M. Hess, Inigo Martincorena and Loris Mularoni for driver analyses using additional methods, Federico Abascal for providing PCAWG germline variant density estimates, Ivan Borozan and Vincent Ferretti for initial analyses of viral genome integration sites, Omar Wagih for kinase binding site analysis, and members of the PCAWG Drivers working group for many useful discussions. The results published here are in part based upon data generated the International Cancer Genome Consortium (ICGC) and The Cancer Genome Atlas (TCGA) Research Network.

